# Adaptation of peristaltic pumps for laminar flow experiments

**DOI:** 10.1101/2021.10.17.464751

**Authors:** Javier Abello, Yvette Y. Yien, Amber N. Stratman

**Author notes:** Correspondence (ANS).

## Abstract

Endothelial cells (ECs) are the primary cellular constituent of blood vessels that are in direct contact with hemodynamic forces over the course of a lifetime. Throughout the body, vessels experience different types of blood flow patterns and rates that alter vascular architecture and cellular behavior. Because of the complexities of studying blood flow in an intact organism, particularly during development, modeling of blood flow *in vitro* has become a powerful technique for studying hemodynamic dependent signaling mechanisms in ECs. While commercial flow systems that recirculate fluids exist, many commercially available pumps are peristaltic and best model pulsatile flow conditions. However, there are many important *in vivo* situations in which ECs experience laminar flow conditions, such as along long, straight stretches of the vasculature. To understand EC function under these situations, it is important to be able to consistently model laminar flow conditions *in vitro*. Here, we outline a method to reliably adapt commercially available peristaltic pumps to reproducibly study laminar flow conditions. Our proof of concept study focuses on 2-dimensional (2D) models but could be further adapted to 3-dimensional (3D) environments to better model *in vivo* scenarios such as organ development. Our studies make significant inroads into solving technical challenges associated with flow modeling, and allow us to conduct functional studies towards understanding the mechanistic role of flow forces on vascular architecture, cellular behavior, and remodeling during a variety of physiological contexts.

## Introduction

Perfusing 2D or 3D cultures of endothelial cells (EC) with fluid is a popular technique to model the effects of blood flow and shear stress forces on the vasculature [1-16]. *In vitro* systems have been built to mimic individual types of flow forces or patterns felt by ECs along different regions of the vascular tree, including oscillatory, pulsatile, and laminar. Consistent with what is observed under physiological conditions, ECs cultured *in vitro* under laminar flow conditions align parallel to the direction of flow [5, 6, 14-29]. Similarly, ECs experiencing pulsatile or oscillatory flow conditions remain more haphazardly organized, or with a perpendicular alignment to the flow direction [5, 6, 14, 15, 17-29].

While significant research has been done to build reliable platforms capable of generating physiologic levels of laminar versus oscillatory/pulsatile flow forces, systems remain inaccessible to the broad vascular biology research community for a number of reasons. For many, the price of a full commercial system capable of altering flow patterns, rates, or flow type prevents use of this modeling technique. Peristaltic pumps are more cost effective but best suited to mimic pulsatory/oscillatory flow conditions, due to the design of the pump controlling the flow rates [30]. To account for this, we have adapted and optimized a commercially available, multi-head peristaltic pump system that has its own controller software [31], to be able to reproducibly perform laminar flow or pulsatile flow experiments (**Fig 1A**). Here, we will outline the steps needed to create pulse dampeners from easy to access supplies, that can attach to a peristaltic pump to generate laminar flow. Further, we discuss our parameters for eliciting flow responses, standardization/validation of this pump system, and downstream analysis techniques for studying cell biology using this platform.

**Fig 1.**
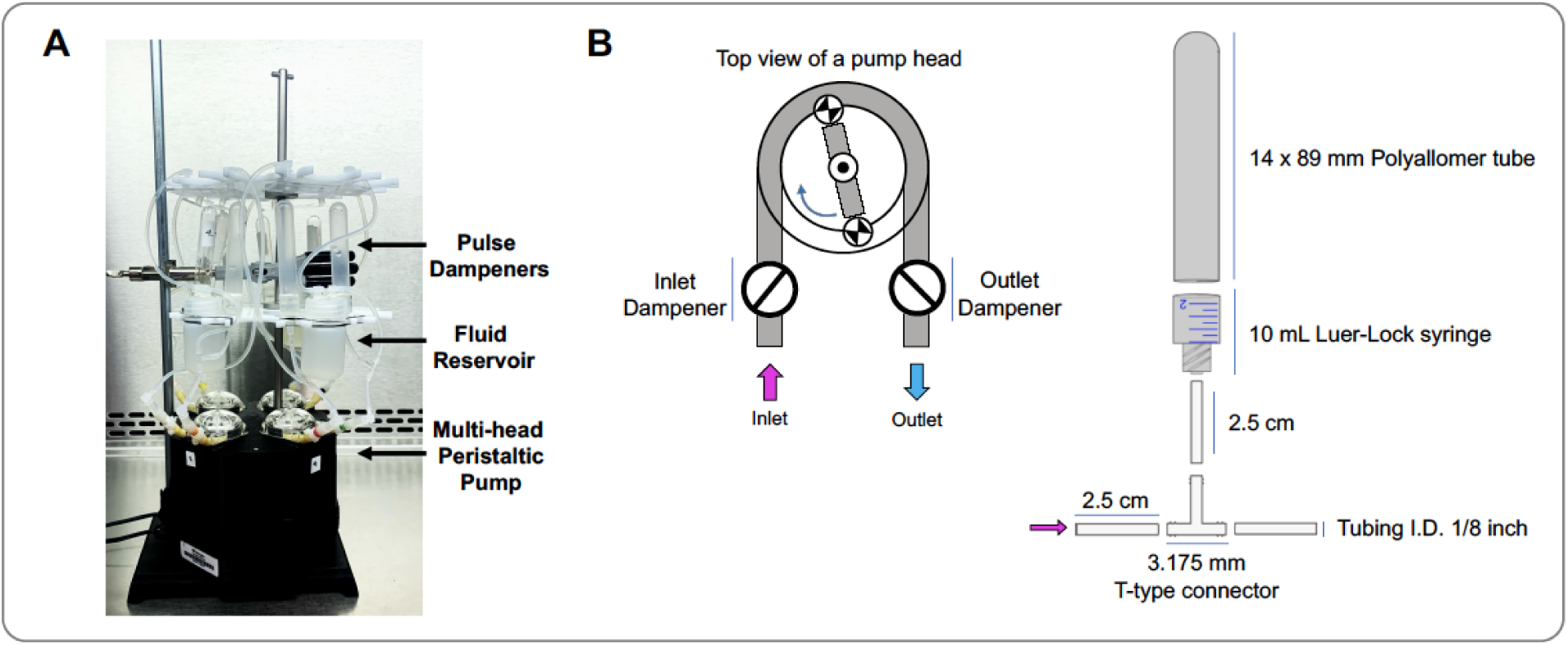
Adaptations to a peristaltic pump to deliver laminar flow. **A)** Picture of the flow system setup, including the 4 head peristaltic pump, dampeners, reservoirs, and slides. **B)** Left panel shows the schematic representation of a peristaltic pump’ head including the positions of the dampeners. The right panel is a schematic representation of how to assemble the dampeners.

## Results

### Generation of dampeners to offset pump pulsation

Addition of a dampener to a peristaltic pump is one way to offset the pulsatile forces generated by the mechanical properties of the pump. To do this, we utilized easy to access and cost-effective supplies to custom make small footprint dampeners to attach to any flow system (**Fig 1B**). To create the body of the dampener, the barrel of a 10 mL syringe (BD) was cut 12 mm from base of the barrel. A thin wall polypropylene tube (Beckman) was bonded to the syringe barrel by applying a thin layer of two-parts urethane adhesive (J-B Weld Company), following the manufacturer recommendation. Because the internal diameter of the syringe barrel is 13.34 mm and the external diameter of the polypropylene tube is 14.43 mm, to assemble the dampener body both parts must be forced together, which along with the adhesive avoids air or liquid leaks. The tubing connection to the dampener was created with 2.5 cm silicone tube (FisherBrand) inserted directly to the syringe Luer-Lok tip. At the other end of the silicone tubing, a T-type connector (Nalgene) was inserted (**Fig 1B**). Two dampeners are connected to the system, one on the inlet and one on the outlet side of the system between the pump head and its respective media reservoir. The dampeners must be vertically oriented to ensure proper function.

The media reservoirs were constructed using 30 mL high density polyethylene (HDPE) bottles (Nalgene). Two holes 4 mm in diameter were drilled in the caps of the HDPE bottles and two elbow luer connectors male (ibidi) were bounded to the caps using the two-parts urethane adhesive described above. At the interior of the cap two pieces of 2 and 4 cm silicone tubing were attached to the elbow connector as inlet and outlet ports respectively.

Efficacy of the generation of laminar flow was confirmed two ways. The first was done via real-time visualization of Quantum dots (Qdots) within the circulating media (**Videos 1 and 2**). As shown in the videos, without the dampener (**Video1**) there is a clear rhythmic stopping of the fluid/Qdots as they transverse the flow chamber (flow is moving left to right across the video). This pulsation is virtually eliminated after addition of dampeners (**Video 2**). To quantify the effectiveness of the dampeners in offsetting the inherent pulsatile nature of the pump, we installed a liquid flow sensor to the silicone tubing (Sensirion) enabling us to determine directionality of flow (positive flow moving left to right across the sensor, negative if flow is moving right to left across the sensor) and the dynamic liquid flow rates of the system (**Fig 2**). The sensor was installed before the inlet attachment point of the μ-Slide | ^0.4^ Luer slide of the flow circuit (**Fig 2A**). The system was evaluated under the following conditions: 1) the flow system without dampeners, using the original function of the pump (pulsatile, **Fig 2A-C**); 2) the flow system modified with dampeners installed at the inlet and outlet of the pump head (laminar, **Fig 2D-F**); and 3) the flow system modified with a commercial dampener (Sensirion) installed before the sensor as is recommended by the manufacturer (laminar, **Fig 2G-I**). The system was placed under our standard experimental conditions (5% CO_2_, 37°C, and 95% humidity for 24 hours), and 15 second measurements of fluid flow taken at 1 and 24 hours. The two time points were chosen to demonstrate the stability of flow forces across the time frame of experimentation. As shown, the non-modified pump as purchased has a rhythmic pulsation to it (**Fig 2B**), that can be dramatically off-set by the incorporation of dampeners into the flow circuit (**Fig 2E**). As a direct comparison, we tested the performance of a commercial dampener, and demonstrate our homemade dampeners were able to out preform a commercial version and suppress pulsation more dramatically (**Fig 2E, H**). While there is still minimal pulsation within the system, it is reduced approximately 6-fold, and reliably allows us to utilize this system to observe phenotypic differences in endothelial cell behavior in response to variation in type of flow stimuli.

**Fig 2.**
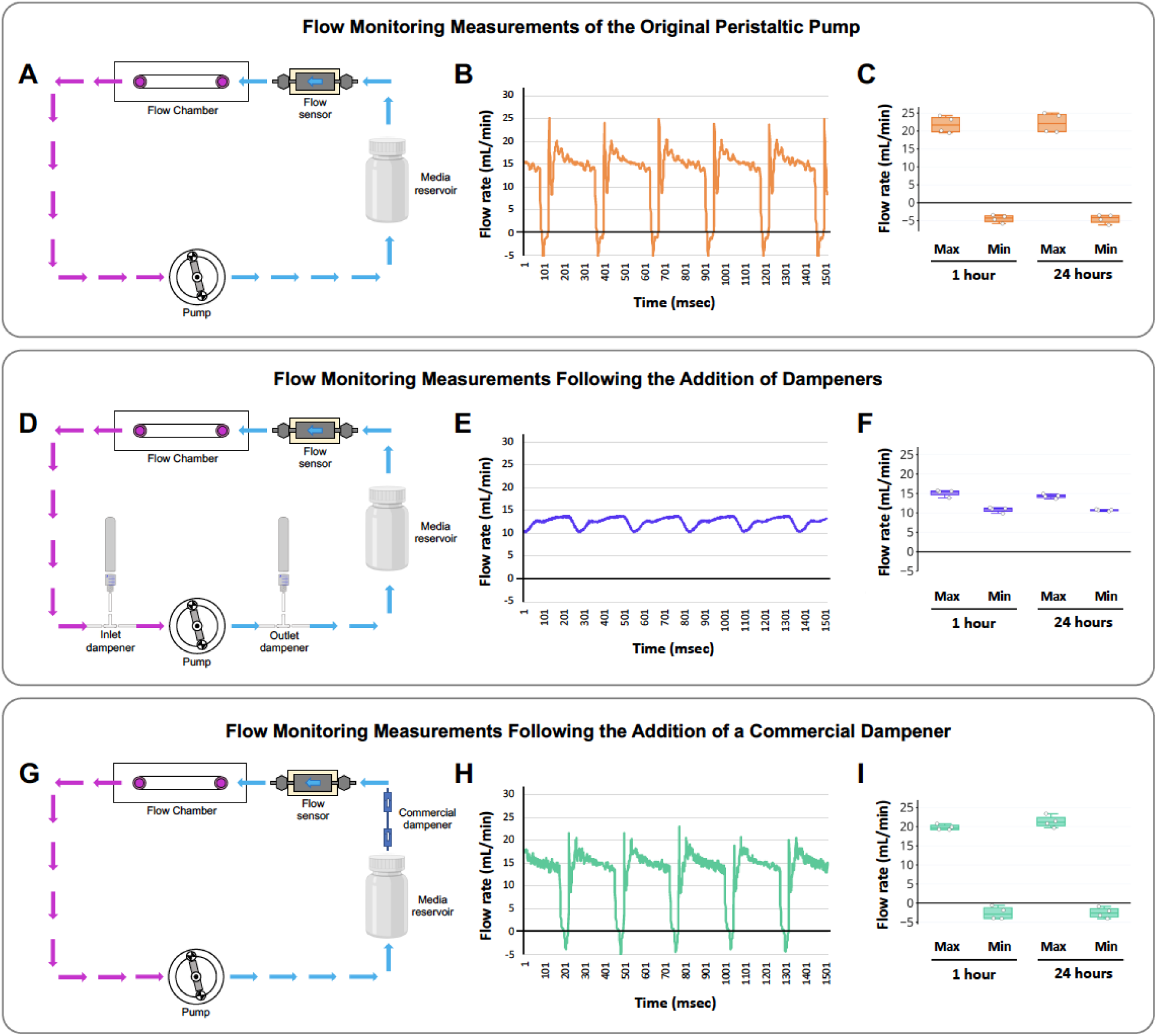
Quantifying the effect of dampeners in off-setting pump pulsation. A flow sensor was purchased to quantifiably measure the effects of dampeners in off-setting peristaltic pump pulsation. The flow sensor was placed directly prior to the flow chamber to ensure a representative measurement of flow forces felt by the endothelial cells in the culture. **A)** Schematic diagram of the flow circuit and placement of the flow sensor to measure flow forces generated by the original, non-modified peristaltic pump. **B)** Pulse traces collected across 1500 msec of pump function, demonstrating marked pulsation of the fluid that is being flowed across the endothelial cell monolayer. **C)** Average maximum and minimum flow forces generated by the endogenous function of the peristaltic pump at 1 and 24 hours of culture. **D)** Schematic diagram of the modified laminar flow circuit, including placement of our custom inlet/outlet dampeners and the flow sensor to measure forces generated by off-setting the pulsation coming out of the peristaltic pump. **E)** Pulse traces collected across 1500 msec of pump function, demonstrating laminar flow (i.e. 6-fold suppression of fluid pulsation) across the endothelial cell monolayer. **F)** Average maximum and minimum flow forces generated by the modified, laminar function of the peristaltic pump at 1 and 24 hours of culture. **G)** Schematic diagram of the modified laminar flow circuit, including placement of a commercial dampener and the flow sensor to measure forces generated by off-setting the pulsation coming out of the peristaltic pump. **H)** Pulse traces collected across 1500 msec of pump function, demonstrating mild suppression of fluid pulsation across the endothelial cell monolayer. However, as shown, our dampeners suppressed pulsation to a much higher degree (commercial 1.4-fold from peristaltic; lab-built 6-fold from peristaltic). **I)** Average maximum and minimum flow forces generated after commercial dampener modifications of the peristaltic pump at 1 and 24 hours of culture. The box plots are graphed showing the median versus the first and third quartiles of the data (the middle, top, and bottom lines of the box respectively). The whiskers demonstrate the spread of data within 1.5x above and below the interquartile range. All data points (averages from individual pumps) are shown as discrete dots, with outliers shown above or below the whiskers.

### Validation of cellular behavior in flow assays via live imaging analysis

To confirm that altered cellular behaviors are generated in response to differing flow stimuli from our modified pump system, we set up 2D live imaging assays to longitudinally track cellular motility and alignment over a 24 hour period. Human umbilical vein endothelial cells (HUVECs) or human aortic endothelial cells (HAECs) were grown to confluence (2.5 × 10^5^ cell/slide) on 1mg/ml gelatin coated μ-Slide | ^0.4^ Luer slides in 1x M199 media supplemented with 20% fetal bovine serum (FBS), 25 μg/mL of Endothelial Cell Growth Supplement (ECGS), 0.01% heparin sodium salt, and 1x Antibiotic-Antimycotic at 5% CO_2_, 37°C, and 95% humidity. The slides were then acclimated to a microscopy system containing a climate-controlled stage top incubator (EVOS M7000) to be cultured under conditions of constant laminar flow, at 12-15 dynes/cm^2^/sec, or constant pulsatile flow, at 12-15 dynes/cm^2^/sec and 60 RPM, for 24 hours. To help maintain cell health, at the start of the experiment the rate of flow was stepwise increased 10% every 10 minutes until reaching the desired final experimental flow rate.

Utilizing a microscope with a stage top incubator allows us to carry out multipoint, time-lapse image acquisition. Images were acquired every 20 minutes for 24 hours to follow dynamic changes in cell shape and alignment over time (**Videos 3 and 4, Fig 3A,B**). At the end of the experiment, the individual images are assembled into a video file to watch cellular behavior across the full 24 hours of imaging. Angle of alignment (**Fig 3C,D**), cellular tracking (**Fig 3E**), and total distance moved/speed of motility (**Fig 3F,G**) amongst other features can be analyzed from these videos utilizing free ImageJ/Fiji software [32]. Our results show that in response to laminar flow, ECs align to and move against the direction of flow over the 24 hour imaging period (**Fig 3D,E, Video 4**), while ECs subjected to pulsatile flow largely remain perpendicular to the direction of flow and move in a more haphazard fashion over the 24 hour imaging period (**Fig 3C,E, Video 3**). Under both flow conditions, HUVECs on average move the same total distance and at the same velocity (**Fig 3F,G**). This behavior mimics what is seen both *in vivo* and *in vitro* flow models [5, 6, 14-29], validating that our system does indeed reliably model flow stimuli to elicit the expected changes in cellular behavior.

**Fig 3.**
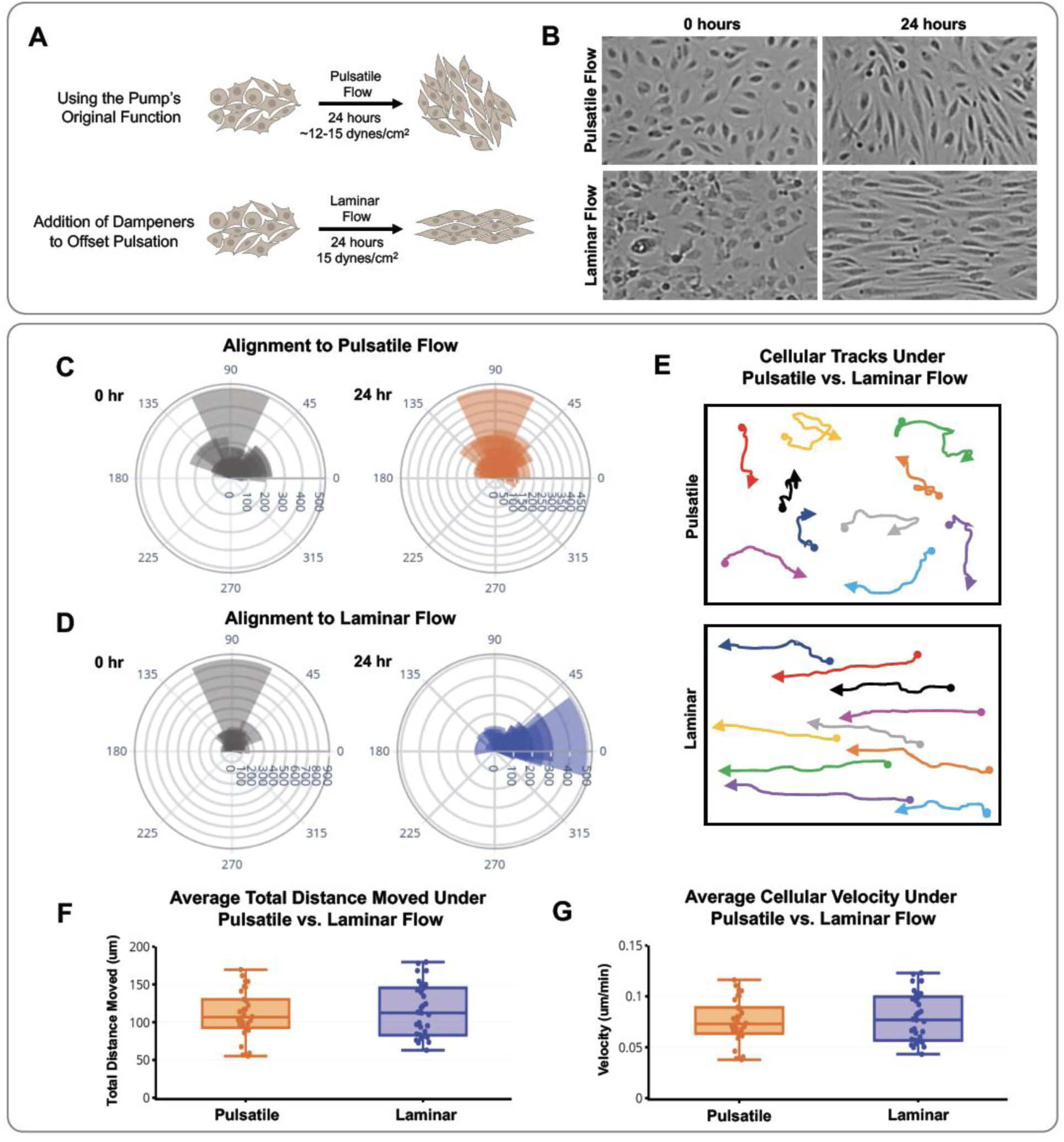
Phenotypic characterization of pulsatile versus laminar flow conditions. **A)** A cartoon of the cell alignment expected under pulsatile and laminar flow conditions (i.e. with and without dampeners). **B)** Microscopy images show HUVEC alignment under pulsatile versus laminar flow conditions at 0 and 24 hours as imaged utilizing our stage top incubated microscope system. **C**,**D)** Distribution angles of HUVECs, showing the orientation of the cell’s longest axis relative to the direction of flow at 0 hours (grey) and after 24 hours (orange/blue) of pulsatile (C) versus laminar (D) flow. Cuboidal or symmetrical cells at 0 hours were measured as 90° to the direction of flow. **E)** Representative cell tracks of HUVECs across the 24 hour period of pulsatile (top) or laminar (bottom) flow. Arrow heads represents the direction of cellular movement, circles represent the starting point of individual cells. **F**,**G)** Average distance traveled (F) and average velocity (G, calculated based off of the average distance travelled over time) shown for pulsatile versus laminar flow over the 24 hour period of treatment. The box plots are graphed showing the median versus the first and third quartiles of the data (the middle, top, and bottom lines of the box respectively). The whiskers demonstrate the spread of data within 1.5x above and below the interquartile range. All data points (individual cells) are shown as discrete dots, with outliers shown above or below the whiskers.

From a practical standpoint, live imaging does not have to be done to utilize this system. If desired, the entire setup can be easily placed in a standard 37°C, 5% CO2, humidified tissue culture incubator. The peristaltic pump we adapted (Flocel [31]) has 4 pressure heads. When doing live imaging, stage top space constraints use of all 4 pumps in tandem. Therefore, we often use the system in a standard incubator to allow conversion of 2 pressure heads to carry laminar flow, while two remain pulsatile to optimize throughput and pair culture conditions within a single experiment. At the end of the experiment, the cells can be collected for biochemical/molecular analysis to determine the downstream molecular impacts of individual flow stimuli: i.e. rinsed with 1x PBS and fixed with 4% paraformaldehyde (PFA) for immunostaining applications (discussed below), flushed with TRIZOL for mRNA collection, or lysed in sample buffer for protein collection [28].

### Immunostaining of cultures following flow

Using our method, following the termination of flow, alterations in protein localization, cellular signaling, or mRNA composition, among other parameters, can be analyzed in order to interrogate downstream signaling pathways that occur as a consequence of flow. Here, we will outline our immunostaining protocol for the analysis of protein accumulation and localization in response to flow forces.

After 24 hours of exposure to flow stimuli, the cultured ECs were rapidly rinsed twice with 3 mL 1x PBS by flushing the slides using a 3 mL syringe. The cells were fixed in 100 μL of 4% paraformaldehyde (PFA) for at least 4 hours before starting the immunostaining protocol. Following fixation, the PFA solution is removed from the slides and the cells rinsed three times with ice cold 1x PBS. If desired, the cultures can then be incubated with 100 μL of 0.1% TritonX-100 in 1x PBS (PBST) for 10 minutes at room temperature, to permeabilize the cells and stain for intracellular proteins. Next, the cells are then incubated in 100 μL blocking buffer (1% BSA, 0.3M glycine in 1x PBST) at room temperature for 1 hour. A primary antibody is chosen for the desired protein of interest, and is diluted into 1% BSA/1x PBST for incubation overnight at 4°C. The next morning, the primary antibody is removed, and the cells rinsed with 1x PBST three times to decrease non-specific antibody binding. Secondary antibody is then added at a 1:2000 dilution in 1% BSA/1x PBST and the slides incubated for 1 hour at room temperature. Finally, HOECHST dye is added at a 1:5000 dilution and incubated for 30 minutes at room temperature for staining of DNA/nuclei. Three washes with PBST are done at the end of all of the steps to remove excess antibody and decrease non-specific background staining prior to mounting the slides for imaging analysis. Images were acquired utilizing a 20x objective on a Nikon Ti2 inverted microscope.

As an example (**Fig 4**), the endothelial cell junctional protein VE-Cadherin was labeled (1:100 dilution of primary antibody) and visualized using an Alexa Fluro-488 secondary antibody (1:2000 dilution, green). Nuclei (HOECHST) labeling is in blue. As shown, application of flow forces across endothelial cells enhances the accumulation of VE-Cadherin protein at junctions compared to no flow controls (**Fig 4A**). While, no differences were noted in immunostaining intensity between laminar and pulsatile flow conditions, marked differences in localization of the VE-Cadherin protein were noted (**Fig 4B-E**).

**Fig 4.**
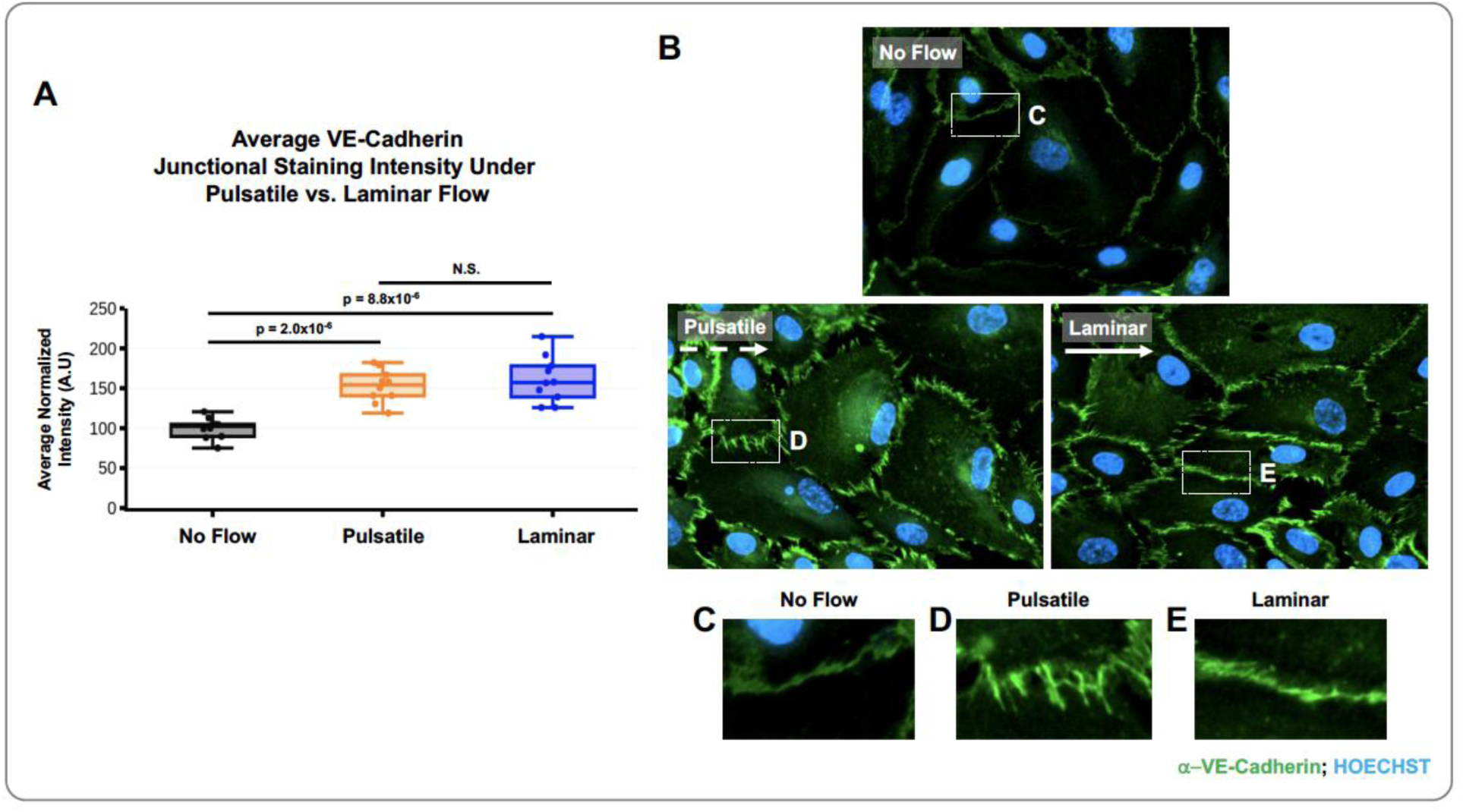
Immunostaining the EC junctional marker, VE-Cadherin, under varying flow conditions. **A)** Quantification of VE-Cadherin staining intensity at the endothelial cell junction. Black bar shows the no flow condition, orange bar shows staining intensity following exposure to pulsatile flow (original pump function), and the blue bar shows staining intensity following exposure to laminar flow (condition with dampeners). The box plots are graphed showing the median versus the first and third quartiles of the data (the middle, top, and bottom lines of the box respectively). The whiskers demonstrate the spread of data within 1.5x above and below the interquartile range. All data points (average intensity from individual images) are shown as individual dots, with outliers shown above or below the whiskers. p-values are indicated above statistically significant datasets and were generated using one-way ANOVA. **B)** Representative microscopy images of cells immunostained for VE-Cadherin (green) and nuclei (blue) under no flow, pulsatile flow, or laminar flow conditions. **C-E)** Zoomed in imaged of the white boxes shown in B, showing changes in localization and intensity of VE-Cadherin staining depending on flow forces applied to the endothelial cell monolayer.

## Discussion

In this work we outline the validation and use of a peristaltic pump system that can be easily modified to concurrently deliver laminar or pulsatile flow to adjacent endothelial cell cultures in a highly reproducible manner. The system provides a cost-effective, easy to use alternative for labs looking to conduct *in vitro* flow modeling experiments. Additionally, the platform allows for real time imaging of cellular responses to flow and analysis of behaviors (**Fig 3**), and following exposure to flow forces, the cells can be utilized for molecular analysis [28] (**Fig 4**).

As part of this work, we compared the endogenous function of the peristaltic pump against our lab-built pulse dampeners and a commercial pulse dampener (**Fig 2**). These studies demonstrated the ability of our lab-built dampeners to generate laminar type flow as compared to the pulsatile nature of the endogenous pump function. Additionally, our dampeners showed improved performance for generation of laminar, non-pulsatile flow forces versus the commercial dampener we tested. The analysis also confirms that our custom-built dampeners are stable across at least 24 hours of assay at the temperature and humidity recommended for endothelial cell culture (**Fig 2C**,**F**,**I**), showing that this system is suitable for use in functional assays. Therefore, modification of a peristaltic pump to include dampeners into the system allows for an affordable and adaptable method to culture cells under laminar and/or pulsatile flow conditions for the reproducible study of shear stress mediated cellular responses *in vitro*.

Functionally, the peristaltic pump utilized has 4 pressure heads, allowing for side by side comparison of cellular behavior under pulsatile versus laminar flow conditions. As shown by cellular tracking experiments, cells under pulsatile flow move haphazardly and tend to end up oriented perpendicular to the direction of flow (**Fig 3A-C,E; Video 3**), while cells under laminar flow rapidly orient parallel to and move against the direction of flow (**Fig 3A,B,D,E, Video 4**). These phenotypes are consistent with those published *in vitro* and *in vivo* under developmental and non-pathogenic flow settings [5, 6, 13-16, 19, 20, 22-27, 29, 31, 33-38], suggesting that this system is able to generate physiologically relevant flow forces. Finally, we confirm that endothelial cells experiencing flow forces develop stronger junctions, as assessed by increased VE-Cadherin localization at junctions compared to no flow conditions (**Fig 4**). While the intensity in staining is not significantly different between cells experiencing pulsatile versus laminar flow, localization of the protein at the junction is differentially oriented (**Fig 4C-E**). Similar results were described previously in ECs cultures, where disturbed flow regions in a flow chamber exhibited discontinuous VE-Cadherin staining, while in areas where the flow was laminar the VE-Cadherin staining was continuous [39].

Being able to reliably model physiological flow conditions allows for deeper mechanistic study of EC autonomous biology; however, it also opens up the possibility of identifying flow regulated signals generated in ECs that alter physiology in a non-autonomous fashion. Embryonic and fetal hematopoietic stem cells (HSCs) and vascular mural cells (MCs) are two primary examples of this phenomenon. HSC development is dependent on blood flow via cell-intrinsic NO signaling [40]. HSC development via the Yes-activated protein (YAP) is also dependent on the mechanical forces of blood flow, as shown by Lundin et al. using induced pluripotent stem cells grown in microfluidic culture devices [41]. Beyond initial specification, HSCs extravasate into circulation and seed developmental niches where they complete their developmental program (reviewed by Horton et al. [42]). During this process, the HSCs are exposed to multiple vascular environments experiencing a variety of mechanical stresses; these stresses trigger signaling events in vascular cells which may play a role in determining ultimate hematopoietic cell fate, among other physiological processes. While there has been tremendous progress in understanding the effects of extracellular forces on HSC differentiation, our understanding of how these forces interface with the vascular niche to signal to HSCs traversing blood vessels has been limited by technical challenges.

Similarly, we have recently shown that differential sensing of blood flow forces can alter mural cell (MC) biology during development [43, 44]. Arteries are known to acquire much greater numbers of MCs than veins across development, and elevated mechanical forces felt by the arterial vasculature is predicted to be a key driver in this process. However, how these forces are sensed by ECs and subsequently communicated to MCs is still an active area of investigation. The transcription factor Klf2 is well known to play an essential role in vascular development in response to forces generated by blood flow [35, 36, 45-53]. In zebrafish and mice, during early vascular development the expression of *klf2a/*Klf2 is significantly higher in veins compared with expression in the dorsal aorta (DA), where the blood flow is pulsatory due to the DAs direct connection to the heart [34, 43, 44]. We recently demonstrated that in *klf2a* deficient zebrafish, there is a significant increase in association of MCs to the CV compared to wildtype siblings, suggesting that Klf2 might serve as a direct or indirect transcriptional repressor for MC recruitment cues [43]. This is just one example of the natural anatomical adaptations that result as a response to the different types of flow forces experienced by vasculature during development or in disease. Therefore, the development of a low cost and easy to use device to model blood flow forces, such as the one described in this manuscript, has the potential to greatly expand our understanding of the physiological effects of EC intrinsic and extrinsic signaling events.

## Materials and Methods

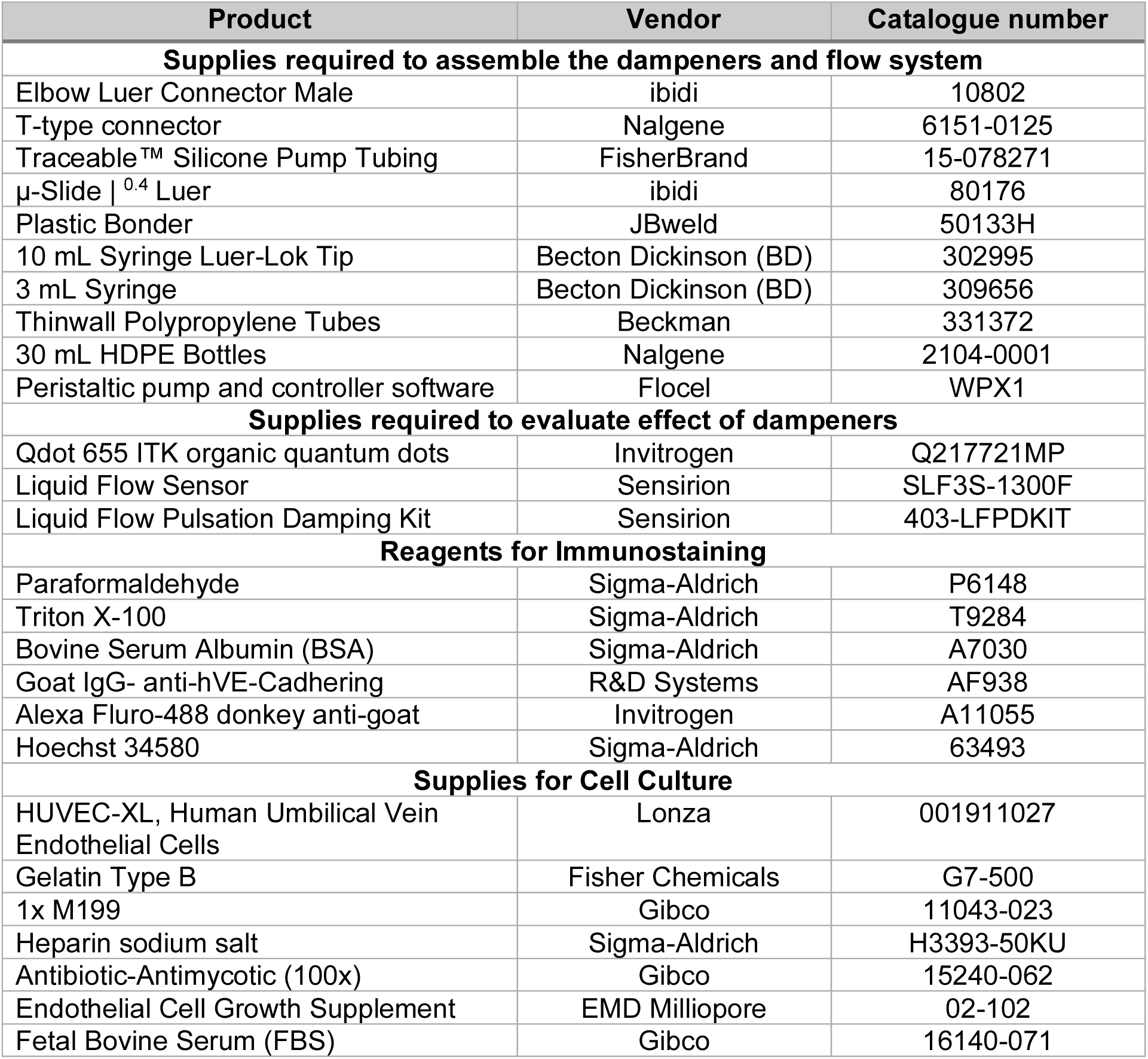

## Acknowledgments

This work was supported by grants from the NIH/NIGMS R35 GM137976 (A.N.S.); Cancer Research Foundation Young Investigator Award (A.N.S.), and NIH/NIGMS R35GM133560 (Y.Y.Y.).

## Supporting information

**Video 1. Visualization of Qdots under pulsatile flow conditions without dampeners**. Qdots were diluted 1:1000 in full culture media and flowed across the imaging slides for visualization in real-time. The video demonstrates Qdot movement under pulsatile flow conditions (the standard pump function without the pulse dampeners present) and a constant pump speed.

**Video 2. Visualization of Qdots under laminar flow conditions with dampeners**. Qdots were diluted 1:1000 in full culture media and flowed across the imaging slides for visualization in real-time. The video demonstrates Qdot movement with dampeners located at the inlet and outlet ports of the pump head to generate laminar flow at a constant pump speed.

**Video 3. Visualization of HUVECs under pulsatile flow conditions (without dampeners)**. Human umbilical vein endothelial cells (HUVECs) were cultured on a μ-Slide | ^0.4^ Luer slide (2.5 × 10^5^ cell/slide) in full culture media. The video shows a 24 hour time lapse sequence, with images acquired at 20 minute intervals. The cells were cultured under pulsatile flow conditions (the standard pump function without the pulse dampeners present) and a constant pump speed.

**Video 4. Visualization of HUVECs under laminar flow conditions (utilizing dampeners)**. Human umbilical vein endothelial cells (HUVECs) were cultured on a μ-Slide | ^0.4^ Luer slide (2.5 × 10^5^ cell/slide) in full culture media. The video shows a 24 hour time lapse sequence, with images acquired at 20 minute intervals. The cells were cultured under laminar flow conditions—i.e. dampeners located at the inlet and outlet ports of the pump head to generate laminar flow at a constant pump speed.

## References

1. Menon NV, Su C, Pang KT, Phua ZJ, Tay HM, Dalan R, et al. Recapitulating atherogenic flow disturbances and vascular inflammation in a perfusable 3D stenosis model. Biofabrication. 2020;12(4):045009. Epub 2020/07/11. doi: 10.1088/1758-5090/aba501. PubMed PMID: 32650321.

2. Terrell JA, Jones CG, Kabandana GKM, Chen C. From cells-on-a-chip to organs-on-a-chip: scaffolding materials for 3D cell culture in microfluidics. J Mater Chem B. 2020;8(31):6667-85. Epub 2020/06/23. doi: 10.1039/d0tb00718h. PubMed PMID: 32567628.

3. Maurya MR, Gupta S, Li JY, Ajami NE, Chen ZB, Shyy JY, et al. Longitudinal shear stress response in human endothelial cells to atheroprone and atheroprotective conditions. Proc Natl Acad Sci U S A. 2021;118(4). Epub 2021/01/21. doi: 10.1073/pnas.2023236118. PubMed PMID: 33468662; PubMed Central PMCID: PMCPMC7848718.

4. Swain SM, Liddle RA. Piezo1 acts upstream of TRPV4 to induce pathological changes in endothelial cells due to shear stress. J Biol Chem. 2021;296:100171. Epub 2020/12/11. doi: 10.1074/jbc.RA120.015059. PubMed PMID: 33298523; PubMed Central PMCID: PMCPMC7948745.

5. Guo D, Chien S, Shyy JY. Regulation of endothelial cell cycle by laminar versus oscillatory flow: distinct modes of interactions of AMP-activated protein kinase and Akt pathways. Circ Res. 2007;100(4):564-71. Epub 2007/02/03. doi: 10.1161/01.RES.0000259561.23876.c5. PubMed PMID: 17272808.

6. Eskin SG, Ives CL, McIntire LV, Navarro LT. Response of cultured endothelial cells to steady flow. Microvasc Res. 1984;28(1):87-94. Epub 1984/07/01. doi: 10.1016/0026-2862(84)90031-1. PubMed PMID: 6748961.

7. Hu YL, Li S, Miao H, Tsou TC, del Pozo MA, Chien S. Roles of microtubule dynamics and small GTPase Rac in endothelial cell migration and lamellipodium formation under flow. J Vasc Res. 2002;39(6):465-76. Epub 2003/02/05. doi: 10.1159/000067202. PubMed PMID: 12566972.

8. LaMack JA, Friedman MH. Individual and combined effects of shear stress magnitude and spatial gradient on endothelial cell gene expression. Am J Physiol Heart Circ Physiol. 2007;293(5):H2853-9. Epub 2007/09/04. doi: 10.1152/ajpheart.00244.2007. PubMed PMID: 17766484.

9. Malek AM, Izumo S. Control of endothelial cell gene expression by flow. J Biomech. 1995;28(12):1515-28. Epub 1995/12/01. doi: 10.1016/0021-9290(95)00099-2. PubMed PMID: 8666591.

10. Shiu YT, Li S, Marganski WA, Usami S, Schwartz MA, Wang YL, et al. Rho mediates the shear-enhancement of endothelial cell migration and traction force generation. Biophys J. 2004;86(4):2558-65. Epub 2004/03/26. doi: 10.1016/S0006-3495(04)74311-8. PubMed PMID: 15041692; PubMed Central PMCID: PMCPMC1304103.

11. Wang S, Tarbell JM. Effect of fluid flow on smooth muscle cells in a 3-dimensional collagen gel model. Arterioscler Thromb Vasc Biol. 2000;20(10):2220-5. Epub 2000/10/14. doi: 10.1161/01.atv.20.10.2220. PubMed PMID: 11031207.

12. Ziegler T, Nerem RM. Effect of flow on the process of endothelial cell division. Arterioscler Thromb. 1994;14(4):636-43. Epub 1994/04/01. doi: 10.1161/01.atv.14.4.636. PubMed PMID: 8148361.

13. Fang JS, Coon BG, Gillis N, Chen Z, Qiu J, Chittenden TW, et al. Shear-induced Notch-Cx37-p27 axis arrests endothelial cell cycle to enable arterial specification. Nat Commun. 2017;8(1):2149. Epub 2017/12/17. doi: 10.1038/s41467-017-01742-7. PubMed PMID: 29247167; PubMed Central PMCID: PMCPMC5732288.

14. Wang C, Lu H, Schwartz MA. A novel in vitro flow system for changing flow direction on endothelial cells. J Biomech. 2012;45(7):1212-8. Epub 2012/03/06. doi: 10.1016/j.jbiomech.2012.01.045. PubMed PMID: 22386042; PubMed Central PMCID: PMCPMC3327813.

15. Miao H, Hu YL, Shiu YT, Yuan S, Zhao Y, Kaunas R, et al. Effects of flow patterns on the localization and expression of VE-cadherin at vascular endothelial cell junctions: in vivo and in vitro investigations. J Vasc Res. 2005;42(1):77-89. Epub 2005/01/08. doi: 10.1159/000083094. PubMed PMID: 15637443.

16. Liu Z, Ruter DL, Quigley K, Tanke NT, Jiang Y, Bautch VL. Single-Cell RNA Sequencing Reveals Endothelial Cell Transcriptome Heterogeneity Under Homeostatic Laminar Flow. Arterioscler Thromb Vasc Biol. 2021;41(10):2575-84. Epub 2021/08/27. doi: 10.1161/ATVBAHA.121.316797. PubMed PMID: 34433297; PubMed Central PMCID: PMCPMC8454496.

17. Vion AC, Perovic T, Petit C, Hollfinger I, Bartels-Klein E, Frampton E, et al. Endothelial Cell Orientation and Polarity Are Controlled by Shear Stress and VEGF Through Distinct Signaling Pathways. Front Physiol. 2020;11:623769. Epub 2021/03/20. doi: 10.3389/fphys.2020.623769. PubMed PMID: 33737879; PubMed Central PMCID: PMCPMC7960671.

18. Steward R, Jr., Tambe D, Hardin CC, Krishnan R, Fredberg JJ. Fluid shear, intercellular stress, and endothelial cell alignment. Am J Physiol Cell Physiol. 2015;308(8):C657-64. Epub 2015/02/06. doi: 10.1152/ajpcell.00363.2014. PubMed PMID: 25652451; PubMed Central PMCID: PMCPMC4398851.

19. Helmlinger G, Geiger RV, Schreck S, Nerem RM. Effects of pulsatile flow on cultured vascular endothelial cell morphology. J Biomech Eng. 1991;113(2):123-31. Epub 1991/05/01. doi: 10.1115/1.2891226. PubMed PMID: 1875686.

20. Dekker RJ, van Soest S, Fontijn RD, Salamanca S, de Groot PG, VanBavel E, et al. Prolonged fluid shear stress induces a distinct set of endothelial cell genes, most specifically lung Kruppel-like factor (KLF2). Blood. 2002;100(5):1689-98. Epub 2002/08/15. doi: 10.1182/blood-2002-01-0046. PubMed PMID: 12176889.

21. Kataoka N, Ujita S, Sato M. Effect of flow direction on the morphological responses of cultured bovine aortic endothelial cells. Med Biol Eng Comput. 1998;36(1):122-8. Epub 1998/06/06. doi: 10.1007/BF02522869. PubMed PMID: 9614760.

22. Mohammed M, Thurgood P, Gilliam C, Nguyen N, Pirogova E, Peter K, et al. Studying the Response of Aortic Endothelial Cells under Pulsatile Flow Using a Compact Microfluidic System. Anal Chem. 2019;91(18):12077-84. Epub 2019/08/14. doi: 10.1021/acs.analchem.9b03247. PubMed PMID: 31407572.

23. Simmers MB, Pryor AW, Blackman BR. Arterial shear stress regulates endothelial cell-directed migration, polarity, and morphology in confluent monolayers. Am J Physiol Heart Circ Physiol. 2007;293(3):H1937-46. Epub 2007/06/26. doi: 10.1152/ajpheart.00534.2007. PubMed PMID: 17586613.

24. Tovar-Lopez F, Thurgood P, Gilliam C, Nguyen N, Pirogova E, Khoshmanesh K, et al. A Microfluidic System for Studying the Effects of Disturbed Flow on Endothelial Cells. Front Bioeng Biotechnol. 2019;7:81. Epub 2019/05/22. doi: 10.3389/fbioe.2019.00081. PubMed PMID: 31111027; PubMed Central PMCID: PMCPMC6499196.

25. Young EW, Simmons CA. Macro- and microscale fluid flow systems for endothelial cell biology. Lab Chip. 2010;10(2):143-60. Epub 2010/01/13. doi: 10.1039/b913390a. PubMed PMID: 20066241.

26. Poduri A, Chang AH, Raftrey B, Rhee S, Van M, Red-Horse K. Endothelial cells respond to the direction of mechanical stimuli through SMAD signaling to regulate coronary artery size. Development. 2017;144(18):3241-52. Epub 2017/08/02. doi: 10.1242/dev.150904. PubMed PMID: 28760815; PubMed Central PMCID: PMCPMC5612251.

27. Flaherty JT, Pierce JE, Ferrans VJ, Patel DJ, Tucker WK, Fry DL. Endothelial nuclear patterns in the canine arterial tree with particular reference to hemodynamic events. Circ Res. 1972;30(1):23-33. Epub 1972/01/01. doi: 10.1161/01.res.30.1.23. PubMed PMID: 5007525.

28. Alghanem AF, Abello J, Maurer JM, Kumar A, Ta CM, Gunasekar SK, et al. The SWELL1-LRRC8 complex regulates endothelial AKT-eNOS signaling and vascular function. Elife. 2021;10. Epub 2021/02/26. doi: 10.7554/eLife.61313. PubMed PMID: 33629656; PubMed Central PMCID: PMCPMC7997661.

29. Coon BG, Baeyens N, Han J, Budatha M, Ross TD, Fang JS, et al. Intramembrane binding of VE-cadherin to VEGFR2 and VEGFR3 assembles the endothelial mechanosensory complex. J Cell Biol. 2015;208(7):975-86. Epub 2015/03/25. doi: 10.1083/jcb.201408103. PubMed PMID: 25800053; PubMed Central PMCID: PMCPMC4384728.

30. Pech S, Richter R, Lienig J, editors. Peristaltic Pump with Continuous Flow and Programmable Flow Pulsation. 2020 IEEE 8th Electronics System-Integration Technology Conference (ESTC); 2020 15-18 Sept. 2020.

31. Cucullo L, Hossain M, Tierney W, Janigro D. A new dynamic in vitro modular capillaries-venules modular system: cerebrovascular physiology in a box. BMC Neurosci. 2013;14:18. Epub 2013/02/08. doi: 10.1186/1471-2202-14-18. PubMed PMID: 23388041; PubMed Central PMCID: PMCPMC3598202.

32. Schindelin J, Arganda-Carreras I, Frise E, Kaynig V, Longair M, Pietzsch T, et al. Fiji: an open-source platform for biological-image analysis. Nat Methods. 2012;9(7):676-82. Epub 2012/06/30. doi: 10.1038/nmeth.2019. PubMed PMID: 22743772; PubMed Central PMCID: PMCPMC3855844.

33. Himburg HA, Dowd SE, Friedman MH. Frequency-dependent response of the vascular endothelium to pulsatile shear stress. Am J Physiol Heart Circ Physiol. 2007;293(1):H645-53. Epub 2007/02/27. doi: 10.1152/ajpheart.01087.2006. PubMed PMID: 17322417.

34. Sugden WW, Meissner R, Aegerter-Wilmsen T, Tsaryk R, Leonard EV, Bussmann J, et al. Endoglin controls blood vessel diameter through endothelial cell shape changes in response to haemodynamic cues. Nat Cell Biol. 2017;19(6):653-65. Epub 2017/05/23. doi: 10.1038/ncb3528. PubMed PMID: 28530658; PubMed Central PMCID: PMCPMC5455977.

35. Boon RA, Leyen TA, Fontijn RD, Fledderus JO, Baggen JM, Volger OL, et al. KLF2-induced actin shear fibers control both alignment to flow and JNK signaling in vascular endothelium. Blood. 2010;115(12):2533-42. Epub 2009/12/25. doi: 10.1182/blood-2009-06-228726. PubMed PMID: 20032497.

36. Chu HR, Sun YC, Gao Y, Guan XM, Yan H, Cui XD, et al. Function of Kruppellike factor 2 in the shear stressinduced cell differentiation of endothelial progenitor cells to endothelial cells. Mol Med Rep. 2019;19(3):1739-46. Epub 2019/01/11. doi: 10.3892/mmr.2019.9819. PubMed PMID: 30628700.

37. dela Paz NG, Walshe TE, Leach LL, Saint-Geniez M, D’Amore PA. Role of shear-stress-induced VEGF expression in endothelial cell survival. J Cell Sci. 2012;125(Pt 4):831-43. Epub 2012/03/09. doi: 10.1242/jcs.084301. PubMed PMID: 22399811; PubMed Central PMCID: PMCPMC3311927.

38. Groenendijk BC, Van der Heiden K, Hierck BP, Poelmann RE. The role of shear stress on ET-1, KLF2, and NOS-3 expression in the developing cardiovascular system of chicken embryos in a venous ligation model. Physiology (Bethesda). 2007;22:380-9. Epub 2007/12/13. doi: 10.1152/physiol.00023.2007. PubMed PMID: 18073411.

39. Chien S. Effects of disturbed flow on endothelial cells. Annals of Biomedical Engineering. 2008;36(4):554–62. doi: 10.1007/s10439-007-9426-3. PubMed PMID: WOS:000254176000006.

40. North TE, Goessling W, Peeters M, Li P, Ceol C, Lord AM, et al. Hematopoietic stem cell development is dependent on blood flow. Cell. 2009;137(4):736-48. Epub 2009/05/20. doi: 10.1016/j.cell.2009.04.023. PubMed PMID: 19450519; PubMed Central PMCID: PMCPMC2722870.

41. Lundin V, Sugden WW, Theodore LN, Sousa PM, Han A, Chou S, et al. YAP Regulates Hematopoietic Stem Cell Formation in Response to the Biomechanical Forces of Blood Flow. Dev Cell. 2020;52(4):446-60 e5. Epub 2020/02/08. doi: 10.1016/j.devcel.2020.01.006. PubMed PMID: 32032546; PubMed Central PMCID: PMCPMC7398148.

42. Horton PD, Dumbali SP, Bhanu KR, Diaz MF, Wenzel PL. Biomechanical Regulation of Hematopoietic Stem Cells in the Developing Embryo. Curr Tissue Microenviron Rep. 2021;2(1):1-15. Epub 2021/05/04. doi: 10.1007/s43152-020-00027-4. PubMed PMID: 33937868; PubMed Central PMCID: PMCPMC8087251.

43. Stratman AN, Burns MC, Farrelly OM, Davis AE, Li W, Pham VN, et al. Chemokine mediated signalling within arteries promotes vascular smooth muscle cell recruitment. Commun Biol. 2020;3(1):734. Epub 2020/12/06. doi: 10.1038/s42003-020-01462-7. PubMed PMID: 33277595; PubMed Central PMCID: PMCPMC7719186.

44. Stratman AN, Pezoa SA, Farrelly OM, Castranova D, Dye LE, 3rd, Butler MG, et al. Interactions between mural cells and endothelial cells stabilize the developing zebrafish dorsal aorta. Development. 2017;144(1):115-27. Epub 2016/12/04. doi: 10.1242/dev.143131. PubMed PMID: 27913637; PubMed Central PMCID: PMCPMC5278630.

45. Atkins GB, Jain MK. Role of Kruppel-like transcription factors in endothelial biology. Circ Res. 2007;100(12):1686-95. Epub 2007/06/23. doi: 10.1161/01.RES.0000267856.00713.0a. PubMed PMID: 17585076.

46. Bhattacharya R, Senbanerjee S, Lin Z, Mir S, Hamik A, Wang P, et al. Inhibition of vascular permeability factor/vascular endothelial growth factor-mediated angiogenesis by the Kruppel-like factor KLF2. J Biol Chem. 2005;280(32):28848-51. Epub 2005/06/28. doi: 10.1074/jbc.C500200200. PubMed PMID: 15980434.

47. Chiplunkar AR, Curtis BC, Eades GL, Kane MS, Fox SJ, Haar JL, et al. The Kruppel-like factor 2 and Kruppel-like factor 4 genes interact to maintain endothelial integrity in mouse embryonic vasculogenesis. BMC Dev Biol. 2013;13:40. Epub 2013/11/23. doi: 10.1186/1471-213X-13-40. PubMed PMID: 24261709; PubMed Central PMCID: PMCPMC4222490.

48. Dekker RJ, van Thienen JV, Rohlena J, de Jager SC, Elderkamp YW, Seppen J, et al. Endothelial KLF2 links local arterial shear stress levels to the expression of vascular tone-regulating genes. Am J Pathol. 2005;167(2):609-18. Epub 2005/07/29. doi: 10.1016/S0002-9440(10)63002-7. PubMed PMID: 16049344; PubMed Central PMCID: PMCPMC1603569.

49. Lee JS, Yu Q, Shin JT, Sebzda E, Bertozzi C, Chen M, et al. Klf2 is an essential regulator of vascular hemodynamic forces in vivo. Dev Cell. 2006;11(6):845-57. Epub 2006/12/05. doi: 10.1016/j.devcel.2006.09.006. PubMed PMID: 17141159.

50. Lin Z, Natesan V, Shi H, Dong F, Kawanami D, Mahabeleshwar GH, et al. Kruppel-like factor 2 regulates endothelial barrier function. Arterioscler Thromb Vasc Biol. 2010;30(10):1952-9. Epub 2010/07/24. doi: 10.1161/ATVBAHA.110.211474. PubMed PMID: 20651277; PubMed Central PMCID: PMCPMC3095948.

51. Parmar KM, Larman HB, Dai G, Zhang Y, Wang ET, Moorthy SN, et al. Integration of flow-dependent endothelial phenotypes by Kruppel-like factor 2. J Clin Invest. 2006;116(1):49-58. Epub 2005/12/13. doi: 10.1172/JCI24787. PubMed PMID: 16341264; PubMed Central PMCID: PMCPMC1307560.

52. Sangwung P, Zhou G, Nayak L, Chan ER, Kumar S, Kang DW, et al. KLF2 and KLF4 control endothelial identity and vascular integrity. JCI Insight. 2017;2(4):e91700. Epub 2017/02/28. doi: 10.1172/jci.insight.91700. PubMed PMID: 28239661; PubMed Central PMCID: PMCPMC5313061 exists.

53. Zhou Z, Tang AT, Wong WY, Bamezai S, Goddard LM, Shenkar R, et al. Cerebral cavernous malformations arise from endothelial gain of MEKK3-KLF2/4 signalling. Nature. 2016;532(7597):122-6. Epub 2016/03/31. doi: 10.1038/nature17178. PubMed PMID: 27027284; PubMed Central PMCID: PMCPMC4864035.

